# Sarcomeric SRX:DRX Equilibrium in Alport and LDLR/P407 Mouse Models of HFpEF

**DOI:** 10.1101/2024.02.20.581314

**Authors:** Ali Kamiar, Monique Williams, Jose M. Capcha, Katarzyna Kazmierczak, Jingsheng Liang, Gary D. Lopaschuk, Keith A Webster, Danuta Szczesna-Cordary, Lina A Shehadeh

## Abstract

Cardiac myosin energetic states that regulate heart contractility define interactions of myosin cross-bridges with actin-containing thin filaments have been functionally linked with the pathology of hypertrophic cardiomyopathy (HCM). In particular, the balance between the disordered relaxed (DRX) and super relaxed (SRX) states that correlate respectively with enhanced force and energy conservation significantly determine myocardial performance and energy utilization. Compelling evidence suggests that a balanced SRX and DRX states proportion is a prerequisite for long-term cardiac health. Whereas roles for altered SRX: DRX proportions in HCM have been studied in depth, the mechanics of sarcomeric dysfunction and SRX: DRX proportions have not been reported in models of acquired heart failure (HF) including HF with preserved ejection fraction (HFpEF). Here, we quantified SRX andDRX myosin populations in two mouse models of HFpEF, including Alport and LDLR/P407 mice that represent cardiorenal/hypertensive and cardiometabolic/hyperlipidemic mouse models of HFpEF, respectively. We report significant changes in the SRX:DRX in both HFpEF mouse models, with an increased DRX state associated with Alport mice and a stabilized SRX state associated with LDLR/P407 mice. These findings correlate respectively with the hypercontractility and metabolic dysregulation with bradycardia phenotypes.

## 1. Introduction

Heart failure with preserved ejection fraction (HFpEF) is an increasingly prevalent syndrome, currently accounting for over 50% of all heart failure (HF) cases^1, 2^. Despite such high prevalence, the pathobiology is poorly understood, at least in part because of the heterogeneous nature of the condition that includes multiple phenogroups with diverse etiologies that converge on a relatively more homogeneous HFpEF clinical presentation^3^. Such etiological diversity hinders both diagnosis and treatment and the utility of animal models to replicate the pathology for mechanistic studies and drug development. The essential features of HFpEF include left ventricle (LV) diastolic stiffening and impaired relaxation, preserved ejection fraction (EF) but depressed systolic reserve, LV hypertrophy, and cardiac fibrosis with comorbidities that can include hypertension, kidney disease, diabetes, obesity, and systemic inflammation^3, 4^. Efficient sarcomere contractility and energy equilibrium are fundamental to cardiac function, and evolving work implicates alterations in the conformation state of filamentous myosin as a possible unifying feature in the etiology of hereditary hypertrophic cardiomyopathy (HCM) and possibly other forms of HF^5^.

Myosin structural conformations regulate force generation and energy expenditure in cardiac and skeletal muscles by fine tuning interactions with thin actin filament. Two myosin conformations defined as disordered relaxed (DRX) state and super relaxed (SRX) state correlate with enhanced force and energy conservation, respectively, such that the proportion of DRX and SRX states determine myocardial performance and energy conservation^6^. The importance of such myosin conformations has been eloquently demonstrated by recent work that assigns disrupted physiological SRX:DRX balance as a root cause of genetic HCM^6–10^. HCM mutations that disrupt the physiological balance of SRX and DRX states alter cardiomyocyte contraction, relaxation, and metabolism and convey increased risks for HF and atrial fibrillation^10^. Targeted modulation of myosin conformation has become a therapeutic strategy for such genetic cardiomyopathies and potentially acquired cardiovascular disease, including HFpEF^6, 11, 12^.

Despite indications that the balance of SRX and DRX conformations are intrinsic to and diagnostic of HF associated with HCM, no such measurements have been reported for human or animal HFpEF, although evidence for disrupted contractile function was recently reported in isolated myofibrils from rodent and porcine models^13^. The possibility that the balance between SRX and DRX represents a unifying feature across phenogroups and potentially a common therapeutic target as it is in HCM prompted us to measure SRX and DRX states in skinned LV muscle fibers from two diverse mouse models of HFpEF representing a cardiorenal phenotype with hypertension secondary to chronic kidney disease (CKD)^14^, and a cardiometabolic/hyperlipidemia phenotype without hypertension, CKD, Type II diabetes (T2D) or obesity^15^.

## 2. Methods

### 2.1. Animals

The data supporting the findings of this study can be obtained from the corresponding author upon reasonable request. All animal procedures were approved by the Institutional Animal Care and Use Committee at the University of Miami, adhering to National Institutes of Health guidelines (IACUC protocol 23–103). Wild type (WT) and *Col4a3*^-/-^ (Alport) mice on 129×1/J background were purchased from Jackson Laboratory and bred in-house.

### 2.2. LDLR/P407 mouse model of HFpEF Development

To develop the model as in our previous work,^15^ ten-week-old WT mice on a 129/J background received a single tail vein injection of 1×10^12^ viral genome sequences (VGS) adeno-associated virus 9–cardiac troponin T–LDLR (AAV9-cTnT-LDLR). This method aimed to selectively direct human LDLR overexpression to the heart. Subsequently, the mice received biweekly intraperitoneal injections of Ploxamer-407 (P407), a selective inhibitor of lipoprotein lipase (LPL) (1g/kg), starting a day after the initial tail injection.

### 2.3. Cardiac function measurements of wild type, Alport and LDLR/P407 mice

Echocardiography, mitral valve (MV) pulse wave, and tissue Doppler were conducted on 8-week-old Alport, 8-week-old wild type, and 4-week-post-treatment LDLR/P407 mice. Cardiac function was assessed using the Vevo2100 imaging system (Visual Sonics, Toronto, ON, Canada) with an MS400 linear array transducer (as in our previous works^16, 17^).

### 2.4. Preparation of skinned Left Ventricular Papillary Muscle (LVPM) fibers

LVPM fibers were prepared as described earlier ^18, 19^. Briefly, muscle bundles were isolated from the hearts of experimental mice (wild type, Alport, and LDLR/P407) and then separated into small muscle strips (2-3 mm in length and 0.5-1 mm in diameter) in ice-cold pCa 8 solution 2.5 mM [Mg-ATP2-], 20 mM MOPS pH 7.0, 15 mM creatine phosphate, and 15 U/ml of phosphocreatine kinase, ionic strength = 150 mM adjusted with KP solution that contained 30 mM 2,3-butanedione monoxime (BDM) and 15% glycerol. The muscle strips were then placed into a pCa 8 solution containing 50% glycerol (storage solution), incubated for one hour on ice, and then immersed in 1% Triton X-100 and 50/50 (%) pCa 8 and glycerol overnight at 4°C. The fibers were transferred into a new storage solution and kept at −20°C until used for experiments.

### 2.5. Determination of proportion of myosin heads occupying the DRX/SRX states

To estimate the number of myosin heads occupying the DRX versus SRX states in wild-type, Alport, and LDLR/P407 mice, Adenosine triphosphate (ATP) turnover measurements were performed using skinned LVPM, as previously described ^18–20^. In these experiments, the fluorescent N-methylanthraniloyl (mant)-ATP was exchanged for nonfluorescent (dark) ATP in skinned LVPM fibers using the IonOptix instrument. First, LVPM fibers were incubated in 250 μM mant-ATP (Thermo Fisher Scientific, Waltham, MA, USA) in a rigor solution [120 mM KPr, 5 mM MgPr, 2.5 mM K_2_HPO_4_, 2.5 mM KH_2_PO_4_, 50 mM MOPS, pH 6.8, and fresh 2 mM DTT] until maximum fluorescence reached a plateau. Then, mant-ATP was chased with 4 mM unlabeled ATP (in rigor buffer), resulting in fluorescence decay. Collected over time, fluorescence intensity isotherms were fitted to a two-exponential decay equation: Y=1-P1[1-exp(-t/T1)]-P2[1-exp(-t/T2)]

Amplitudes of the fast (P1) and slow (P2) phases of fluorescence decay and their respective T1 and T2 lifetimes (in seconds) were derived from fits to a nonlinear least-squares algorithm in GraphPad PRISM version 10 (GraphPad Software, San Diego, CA, USA) ^18–20^. T1 and T2 are the lifetimes of DRX and SRX states. The numbers are derived from the fits to double exponential equation. The P1 was corrected for the fast release of nonspecifically bound mant-ATP in the sample and was derived experimentally using a competition assay, as described in ^18, 21^. The fraction of nonspecifically bound mant-ATP in LVPM fibers was equal to 0.44±0.02, and the population of myosin heads directly occupying the SRX state was then calculated as P2/(1-0.44). All studies were conducted at room temperature (∼23°C).

### 2.6. Statistical Analysis

One-way ANOVA with the Tukey multiple comparison test was employed for all experiments involving three groups. A significance level of P < 0.05 was utilized, and all tests were two-sided. The data are presented as means ± SD. GraphPad Prism 10 software was employed for all analyses and graph generation.

## 3. Results

### 3.1. Diastolic function of Alport and LDLR/P407 HFpEF mice

MV pulse wave and tissue Doppler echocardiography of 8-week-old Alport, and4-week-post-treatment LDLR/P407 mice versus untreated WT mice revealed similar diastolic dysfunction of both HFpEF models Treated mice displayed preserved EFs (Fig. 1A), prolonged isovolumic relaxation (IVRT) (Fig. 1B) and MV deceleration (MV Decel) times (Fig. 1C). MV E/Eʹ showed an increased trend in both models (Fig. 1E), whereas MV E/A was significantly reduced in Alport mice, but not in the LDLR/P407 group. Representative images of mitral valve pulse wave Doppler and tissue Doppler are shown for each group (Fig. 1F). We previously reported significantly increased cardiac fibrosis, cardiac myocyte area, exercise intolerance, and heart weight/ body weight (HW/BW) in mice from both treatment groups, moderate hypertension in Alport but not LDLR/P407 mice, and markedly reduced lifespan of 10-12 weeks in mice from both treatment groups ^15, 17, 22^.

**Figure 1.**
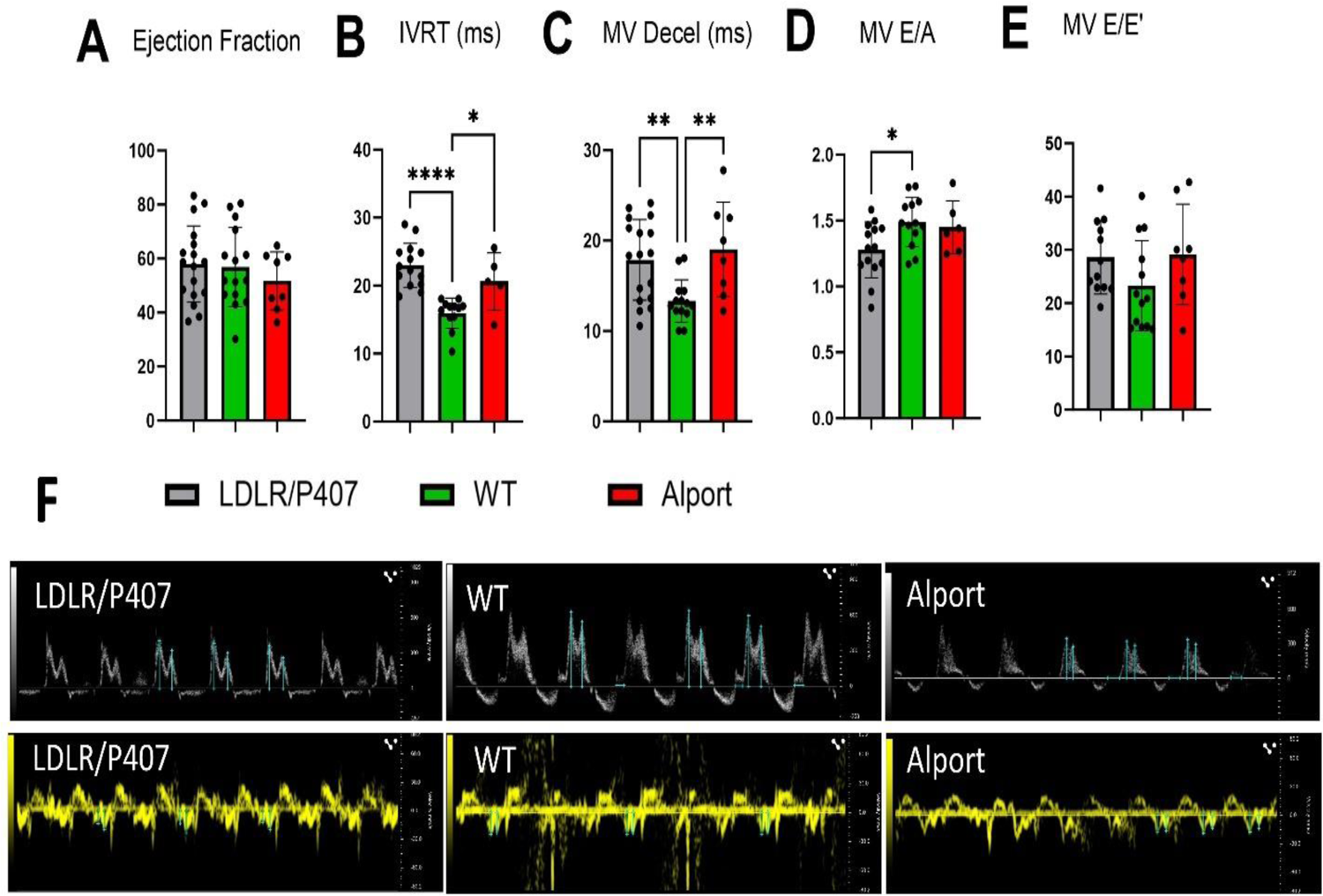
Diastolic function parameters (echocardiography) in WT, LDLR/P407 and Alport mice A. Ejection fraction B. Isovolumic relaxation time (IVRT). C. Mitral valve deceleration time. D. Mitral valve ratio of peak velocity flow in early diastole (the E wave) to peak velocity flow in late diastole (the A wave). E. Mitral valve ratio of peak velocity flow in early diastole (the E wave) to early diastolic mitral annulus velocity (the E’ wave). F. Representative images of pulse wave doppler and tissue doppler. Data are presented as means ± SD. N = 5-17 per group. Statistical significance was determined using one-way ANOVA with Tukey’s multiple comparisons.

### 3.2. The SRX State of Myosin is affected differently in two Mouse Models of HFpEF

Proportional contents of SRX and DRX myosin conformations in skinned LVPM fibers from 8-week-old Alport mice, 8-week-post treated LDLR/P407 mice, and age-matched WT controls are shown in Figure 2 A and B. The content of myosin cross-bridges occupying the SRX state in LVPM fibers of LDLR/P407 mice was significantly greater than that of controls. In contrast, the Alport mouse fibers contained significantly more DRX state myosins, such that LDLR/P407 LVPM contained >10% more SRX state myosins relative to Alport LVPM. Conversely, Alport LVPM contained ∼12% more DRX state myosins (see Fig. 2 pie charts and Table 1). Table 1, in Figure 2B also shows the proportions and lifetimes of the DRX (T1) and SRX (T2) states. T1 was shorter in LDLR/P407 hearts compared to the WT control group (p=0.0397).

**Figure 2.**
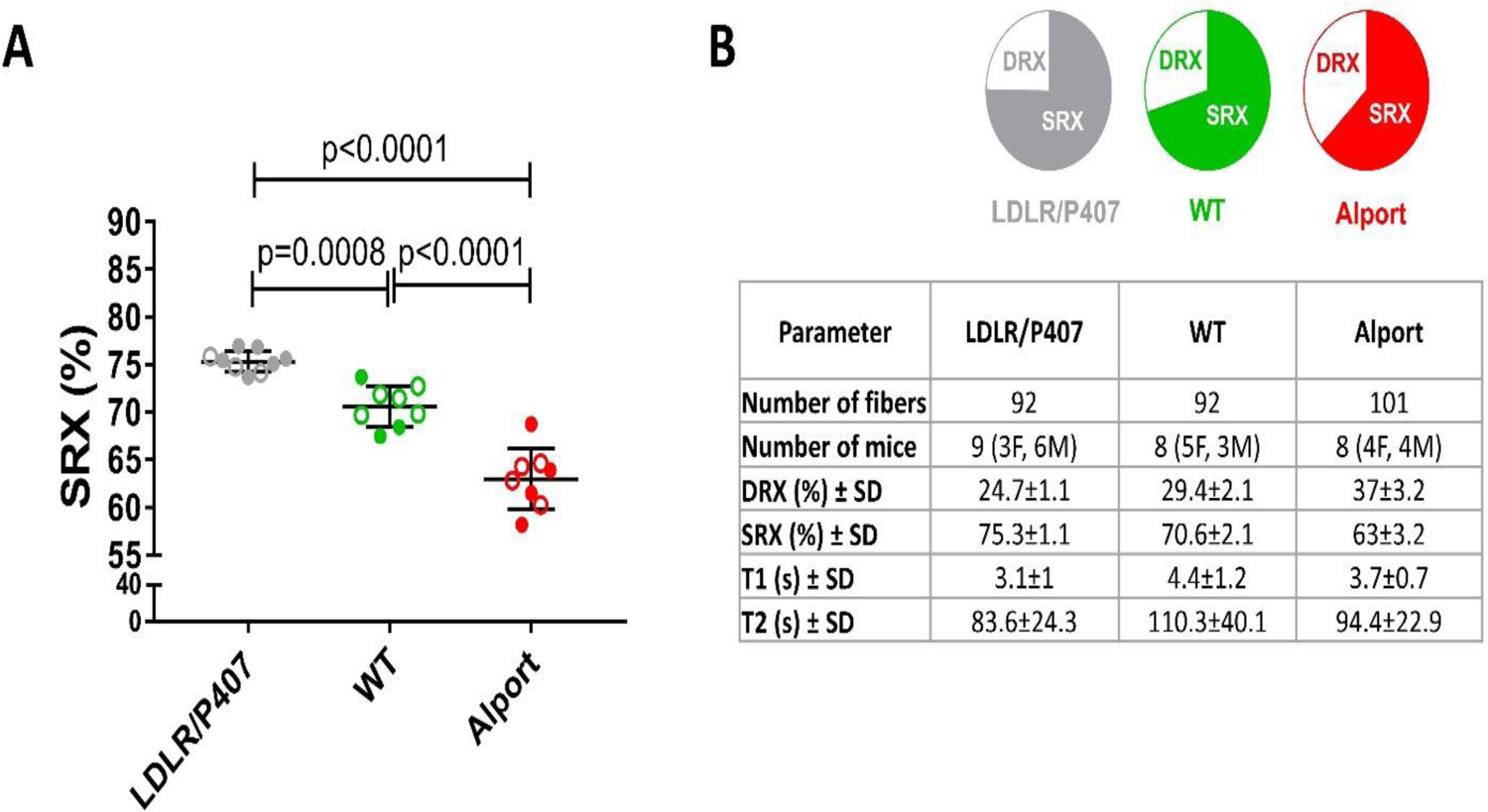
Study of the super-relaxed (SRX) state of myosin in LVPM fibers from WT, Alport, and LDLR/P407 mice. A. Percentage of myosin heads in the SRX state, with closed and open circles indicating male and female mice, respectively. Data are presented as means ± SD. N = 8-9 mice per group. Statistical significance was determined using one-way ANOVA with Tukey’s multiple comparisons. B. Distribution of myosin heads between the SRX and DRX states, illustrated in pie charts. The table includes the lifetimes (in seconds) of the DRX (T1) and SRX (T2) states.

## 4. Discussion

We demonstrate significant changes in the myofibrillar contents of SRX and DRX myosin conformations in two HFpEF mouse models, but unexpectedly, the shifts were in different directions with increased SRX state in LDLR/P407 fibrils and increased DRX state in Alport fibrils. In WT *129J* mice, the proportion of SRX: DRX was 70.6%:29.4%, generating a ratio of 2.4 compared with 75.3%:24.7%, ratio of 3.0 in LDLR/P408 fibrils and 63.0%:37.0%, ratio of 1.7 in Alport fibrils. The ratios on the *129J* background are higher than the theoretical SRX and DRX proportion of 1.5 predicted for normal cardiac muscle^7, 9^. LDLR/P407 fibrils had 4.7% and 12.3% more SRX state myosin respectively than WT and Alport fibrils. The observed changes are physiologically relevant. Changes in myofibrillar SRX:DRX proportions of <10% caused by gain of function HCM mutations in *MYH7* or *MYBPC3* genes expressed in 129SvEv mice were shown to be reversible by mavacamten and significantly responsible for the HCM pathology^6–10^. It is worth mentioning that despite the different patterns of SRX:DRX equilibrium as well as different etiologies in two mouse models, the echo parameters indicate similar levels of diastolic dysfunction.

In another example, the myosin SRX state content of LV myofibers in hibernating ground squirrels increased from 65% during summertime arousal or interbout euthermia to 75% during hibernating torpor, a state of deficient cardiac demand and limited energy resources^6^. Such reapportioning indicates that cardiac SRX:DRX proportions are flexible, in dynamic equilibrium with and regulated by ambient cardiac demands, metabolic intermediates, and energy substrates.^6^ Physiological regulators of SRX:DRX proportions include metabolic intermediates and inotropic effectors, Ca^2+^, heart rate, stretch, and β-adrenergic stimulation that determine the phosphorylation of regulatory contractile proteins.^23^ Evidence suggests a balanced SRX:DRX proportion is a prerequisite for long-term cardiac health ^5–8, 12^.

In the SRX state, the myosin heads are presumed to interact with one another and with the thick filament backbone.^6^ The myosin motor domains are sequestered away from the thin filament, supporting a quiescent, relaxed, energy-conserving state ^8, 21^. In DRX state, more myosin cross-bridges are recruited, and they are readily available to interact with thin filaments and produce force, consuming 5-fold more ATP than the cross-bridges occupying the SRX state^21^. Destabilization of the SRX state in favor of DRX state promotes contractile abnormalities, morphological and metabolic remodeling, and adverse outcomes in animal models and patients ^6, 10^. All causal human HCM mutations tested thus far in mouse and/or in vitro models conferred increased DRX, such that strategies to reinstate intact SRX conformations are being developed as potential treatments for genetic HCM ^11, 12^. Because HCM and HFpEF share common traits that include hypertrophy, impaired relaxation, cardiac fibrosis, HF, arrhythmias, and sudden cardiac death, we expected to find decreased SRX:DRX myosin in both HFpEF models. However, DRX state myosin increased only in Alport mice, whereas fibrils from LDLR/P407 hearts acquired ∼5% more SRX state myosin, a state more reminiscent of energy-conserving torpor squirrel myofibrils than those of hypercontractile cardiac myocytes with HCM mutations (Fig. 2).

At the time of harvest, LV diastolic dysfunction and hypertrophy were well established in both models, as were the HFpEF sequelae of advanced exercise intolerance and imminent sudden death, consistent with energy compromise and/or arrhythmia^14, 15, 17, 22^. Increased DRX state proportion associated with HCM confer energetic and metabolic stresses that include increases of mitochondrial oxygen consumption, citric acid cycle flux, nicotinamide adenine dinucleotide: Reduced Nicotinamide Adenine Dinucleotide (NAD: NADH), and glycolysis, as well as depressed phosphocreatine, and all intermediates of hypercontractility, and reversible by pharmacological rebalancing to physiological SRX:DRX ^6^. In our LDLR/P407 cardiometabolic model, inhibition of LPL prevents cardiac uptake of fatty acid from circulating Very-low-density-lipoproteins (VLDLs), depriving the heart of a major source of fuel, and potentially promoting an energy deficit and metabolic stress^15^. The LDLR/P407 mouse model also exhibit atrioventricular heart blocks^15^ and bradycardia, a common sequela of HFpEF indicative of a low energy state^24^. These factors share a commonality with the torpor squirrel and may be more conducive to an increased SRX:DRX proportion. Decreased cardiac triglyceride (TG) and fatty acid (FA) metabolism have been causally linked with contractile dysfunction, hypertrophy, perivascular fibrosis, and HF^25–29^.

Using isolated myofibers, Fenwick *et al.* recently reported reduced relaxation rates in two rodent HFpEF models, depressed systolic myofibrillar function in a pig model, and no change of fiber resting tension in any model^13^. All models mimicked a hypertensive, cardiometabolic/obesity phenogroup. The authors concluded that whereas myofibril mechanics may uniquely recapitulate distinct aspects of some human HFpEF subphenotypes, changes in addition to those at the sarcomere level including other cellular- and tissue-level pathologies are required to account for the global changes in diastolic dysfunction of HFpEF. From our results we conclude that a decline of the SRX:DRX proportion is not a prerequisite for HF in these models. However, the presence of significant shifts of the SRX:DRX proportion in both models, albeit in opposite directions, is consistent with the involvement or equilibrium of such structure/function changes at the sarcomere level of the sarcomere with the HFpEF phenotype. Further studies on additional HFpEF phenogroup models, including large animals, are warranted.

## 5. Conclusion

Myofibers from Alport mice that model a hypertensive, cardiorenal HFpEF phenogroup contained 7.6% more DRX state myosins than fibrils from the age-matched WT *129J* background strain, consistent with a hypercontractile state and disrupted SRX:DRX proportion that may contribute to the HFpEF phenotype. Conversely, fibrils from LDLR/P407 mice that model a cardiometabolic/hyperlipidemic phenogroup contained 4.7% more SRX state myosins, suggesting a more relaxed, low cardiac output energy conserving state. LDLR/P407 mice contain significantly depressed cardiac TG compared with WT mice, display heart blocks^15^ and bradycardia, and have severe exercise intolerance^15^, suggesting a low energy, hypocontractile state perhaps reminiscent of torpor squirrel sarcomeres that may contribute to increased SRX:DRX proportion, although the latter condition is much more extreme. Respiratory, and metabolic rates of the 13-lined ground squirrel fall to 3% of basal levels during torpor, and HR falls from ≈340 bpm during arousal to ≈6 bpm in torpor^30^.

## Acknowledgement

We thank the Penncore and NHLBI Gene therapy Resource Program (GTRP) for making the Adeno-Associated Viruses used in this project.

## 6. Sources of Funding

Funded by grants from the National Institute of Health (NIH) (1R01HL140468; LAS) and the Miami Heart Research Institute to LAS, R01-HL143830 to DSC and R01 EY033805 to KAW. MW was a recipient of NIH Diversity Supplement Award from 2020 – 2022 (R01HL140468-03S1).

## 7. Disclosures

The authors have no conflicts to disclose.

## Non-standard Abbreviations and Acronyms

AAV9-cTnT-LDLR: Adeno-associatedvirus9-cardiacTroponinT-LDL Receptor

IVRT: Isovolumic relaxation time

P407: Poloxamer-407

HFpEF: Heart failure with preserved ejection fraction

LV: Left ventricle

EF: Ejection fraction

HF: Heart failure

HCM: Hypertrophic cardiomyopathy

DRX: Disordered relaxed state

SRX: Super relaxed state

CDK: Chronic kidney disease

T2D: Type II diabetes

WT: Wild type

VGS: Viral genome sequences

LPL: Lipoprotein lipase

MV: Mitral valve

LVPM: Left Ventricular Papillary Muscle

(ATP): Adenosine triphosphate

MV: Decel Mitral valve deceleration

TG: Triglyceride

FA: Fatty acid

VLDLs: Very-low-density-lipoproteins

HW:BW: Heart weight: Body weight

NAD: NADH Nicotinamide adenine dinucleotide: Reduced Nicotinamide Adenine Dinucleotide

## Notes

### Competing Interest Statement

The authors have declared no competing interest.

